# Drift-Diffusion Modeling Reveals Dissociable Components of Perceptual Learning

**DOI:** 10.1101/2025.09.17.676811

**Authors:** Nikan Amirkhani, Mehdi Sanayei

## Abstract

Visual perceptual learning leads to improved accuracy and faster reaction times. This effect is specific to the location and lower-level features of the training stimulus, as well as other task parameters. However, extensive practice using a specific set of stimulus conditions has been shown to partially generalize to new conditions in the form of faster learning rates and slightly improved performance at the outset. We hypothesized that this effect is caused by nuanced changes in the subject’s decision-making strategy that can be revealed through drift-diffusion modeling (DDM). To test this hypothesis, subjects were trained on a coarse direction discrimination task using random dot motion stimuli over several sessions. Stimulus location and direction of motion were fixed during training, and either of them changed on the testing session. A subset of participants were trained on separate days to test for the effect of sleep consolidation. We observed the typical training effects of improved accuracy and reaction times as a function of training. DDM modeling showed that learning induced increased drift rate, decreased boundary height, lowered integration leak, and decreased rate of the collapsing bound. Importantly, while changes to the drift rate and boundary conditions mostly reverted to their pre-training values on the testing session, integration leak and rate of the collapsing bound remained at their post-training values in the new stimulus conditions. Sleep consolidation exerted no meaningful difference on performance, reaction times, or parameter values. These results suggest that the observed changes to accuracy and reaction times are caused by at least two different components with different degrees of transferability, as well as the fruitfulness of DDM for understanding the nuances of perceptual learning.

## Introduction

Perceptual learning is the improvement of perceptual skills through practice. Seminal studies provided clear demonstrations that performance on visual tasks, such as vernier acuity, could improve dramatically with training (Mckee & Westhe, 1978). Early research showed the specificity of perceptual learning to basic visual stimulus features, such as location (Fahle & Morgan, 1996), orientation (Fahle & Edelman, 1993; Fiorentini & Berardi, 1980), and direction of motion (Ball & Sekuler, 1982). As summarized in an early review (Sagi & Tanne, 1994), when trained stimuli were presented at a new location or with an orthogonal orientation, performance often deteriorated to pre-training levels. This specificity underpinned the influential hypothesis that perceptual learning primarily involves plastic changes in early visual areas, where neurons are finely tuned to such attributes (Hubel & Wiesel, 1962; Tootell et al., 1988). However, subsequent psychophysical work revealed that this specificity is not absolute. For instance, learning was found to transfer across different orientations, leading to the proposal that observers may learn a “rule” for the task rather than a specific stimulus template (Saarinen & Levi, 1995). More strikingly, the classic constraint of location specificity could be entirely overcome with a “double training” paradigm, suggesting that under certain training regimes, higher-level processes may be engaged in perceptual learning and can enable location transfer (Xiao et al., 2008).

These findings fueled a critical debate: does learning reflect a true change in early sensory representations, or the adoption of more effective cognitive strategies? Some argued that many instances of “learning” might not reflect low-level plasticity at all, but rather the observer learning to attend to the most relevant information or apply a better decision-making rule (Mollon & Danilova, 1996). To reconcile these views, the “Reverse Hierarchy Theory” was proposed, suggesting that learning begins with modifications to high-level, decision-related areas and only proceeds to refine low-level sensory representations when the task is sufficiently difficult and demands higher resolution (Ahissar & Hochstein, 2004). This framework provided a compelling theoretical account for why plasticity might be observed at different levels of the visual system.

The search to identify the neural locus of learning has attempted to resolve this debate. Following the logic of the early psychophysical work, initial hypotheses focused on the primary visual cortex (V1) (Schoups et al., 2001). However, electrophysiological studies in non-human primates challenged this view. Contrary to expectations of robust V1 plasticity, training produced at most subtle changes in V1, which were largely attributed to top-down feedback from higher areas (Yang & Maunsell, 2004; Sanayei et al., 2018). Instead, plasticity was more consistently observed in mid-level ventral areas such as V4, whereas dorsal areas at the same level showed no training-related changes. The strongest evidence came from Law and Gold (Law & Gold, 2008), who demonstrated that improved motion discrimination was associated with changes in lateral intraparietal cortex (LIP), rather than in middle temporal cortex, suggesting that learning occurs at the decision-making stage rather than early sensory encoding.

As researchers used fMRI to investigate visual perceptual learning in human subjects, with tasks similar to ones that were used in non-human primates a decade earlier, a different picture emerged. These studies show changes not only in cortical areas involved in decision making, including the LIP, ventral premotor cortex, and the frontal eye field (Chen et al., 2015; Jia et al., 2018; Shibata et al., 2016), but also in early and mid-level visual areas. For instance, training on orientation discrimination was linked to modulation in V1 and V2 (Chen et al., 2016; Mukai et al., 2007), while tasks involving texture and shape discrimination engaged higher-tier regions such as V4 and the lateral occipital complex (Maertens & Pollmann, 2005). These findings suggest that perceptual learning may involve both sensory plasticity and changes in how sensory evidence is integrated in decision-making.

Beyond the locus of neural changes, another important factor shaping perceptual learning is consolidation. A large body of work has shown that improvements in visual discrimination can depend critically on post-training consolidation, with sleep often facilitating performance gains that do not emerge during wakefulness (Gais et al., 2000; Karni et al., 1994; Stickgold et al., 2000). However, there is some evidence that the benefits of sleep for learning may be task-dependent, with Doyon et al. seeing no evidence of added performance gain in a visuomotor adaptation task (Doyon et al., 2009). These findings suggest that consolidation effects are not simply a function of repeated exposure, but are also governed by offline processes that stabilize and reorganize neural representations in a task-specific manner.

A still open question is whether perceptual learning is primarily stimulus-dependent (reflecting changes in early sensory representations) or stimulus-independent (reflecting modifications in decision-related processes). To address this question, we drew on the perceptual decision-making literature, which over the past three decades has used random-dot motion (RDM) tasks and computational modeling to identify neural correlates of evidence accumulation and choice (Gold & Shadlen, 2007). In particular, the drift diffusion model (DDM) has proven powerful for linking behavioral improvements to underlying cognitive mechanisms, and has previously been applied to perceptual learning in orientation discrimination tasks (Petrov et al., 2011). Here we extend that approach to motion discrimination by using a more generalized DDM (Shinn et al., 2020) that allows for a richer characterization of parameters governing accuracy and reaction time across training. This framework enables us to test how different computational components of decision-making contribute to learning.

In this study, we trained participants on a free-response version of the classic RDM task in multiple sessions. In the last session, we tested the generalization of training to a new location or axis of motion in human participants. In another experiment, the training sessions were spaced across multiple days to examine the role of sleep-dependent consolidation in learning and transfer. We fitted the DDM to reaction time and performance on each session to obtain parameters related to sensory (drift rate) and decision (boundary height, drift leak, boundary collapse rate) to tease apart the contribution of each of these parameters to learning. We found that all parameters changed through training, but the pattern of generalization was different between them. While drift rate and boundary height were only partially generalized to new stimulus features, drift leak and boundary collapse rate were fully generalized. Furthermore, these results were not dependent on sleep. These results show that both stimulus-dependent and stimulus-independent (or decision-related) mechanisms may play a crucial role in perceptual learning.

## Methods

### Apparatus and stimuli

Testing occurred in a dimly lit room where participants were seated with their chin and forehead stabilized by a chin and head rest, ∼60cm from a 17″ (1920 ×1080 resolution) monitor with a 60 Hz refresh rate. Experiments were run using a PC running Linux OS. Stimulus presentation and data collection were done using MATLAB (R2016a) and Psychtoolbox (Brainard, 1997; Kleiner et al., 2007; Pelli, 1997). The position of the left eye was monitored throughout the experiment (EyeLink 1000, SR Research) at 1KHz. The stimulus was a random dot motion (RDM) stimulus, presented within a 5° aperture (diameter) on a gray background (RGB = 50). Each dot (white, RGB = 255) subtended 2 pixels on each side and moved at a speed of 5°/s. The density of the dots was 16.7 dots/degree^2^/s. These parameters were the same as those of most decision-making studies that used RDM (Palmer et al., 2005).

### Task Structure and Protocol

Participants engaged in a four-session training regimen followed by a single test session (**Figure 1**) within a single day (in Experiments 1 and 2). There were two forms of trials: (1) Free response (FR) trials in which subjects made a saccade to a pre-determined choice target in a two-alternative forced-choice design, whenever they reached a decision, and (2) Speeded decision (SD) trials where subjects are instructed to respond as quickly as possible while maintaining high accuracy. Each session comprised four FR blocks, each containing 110 trials and one SD block with 20 trials. Each trial was preceded by a grey fixation cross for FR and a white fixation cross for SD blocks. In FR trials, dots moved with coherence levels from 0.0, 3.2, 6.4, 12.8, 25.6, and 51.2%, chosen pseudo-randomly on each trial. The coherence of dots in the SD trials was fixed at 100%. In total, each session consisted of 440 FR trials (40 trials per coherence) and 20 SD trials.

**Figure 1:**
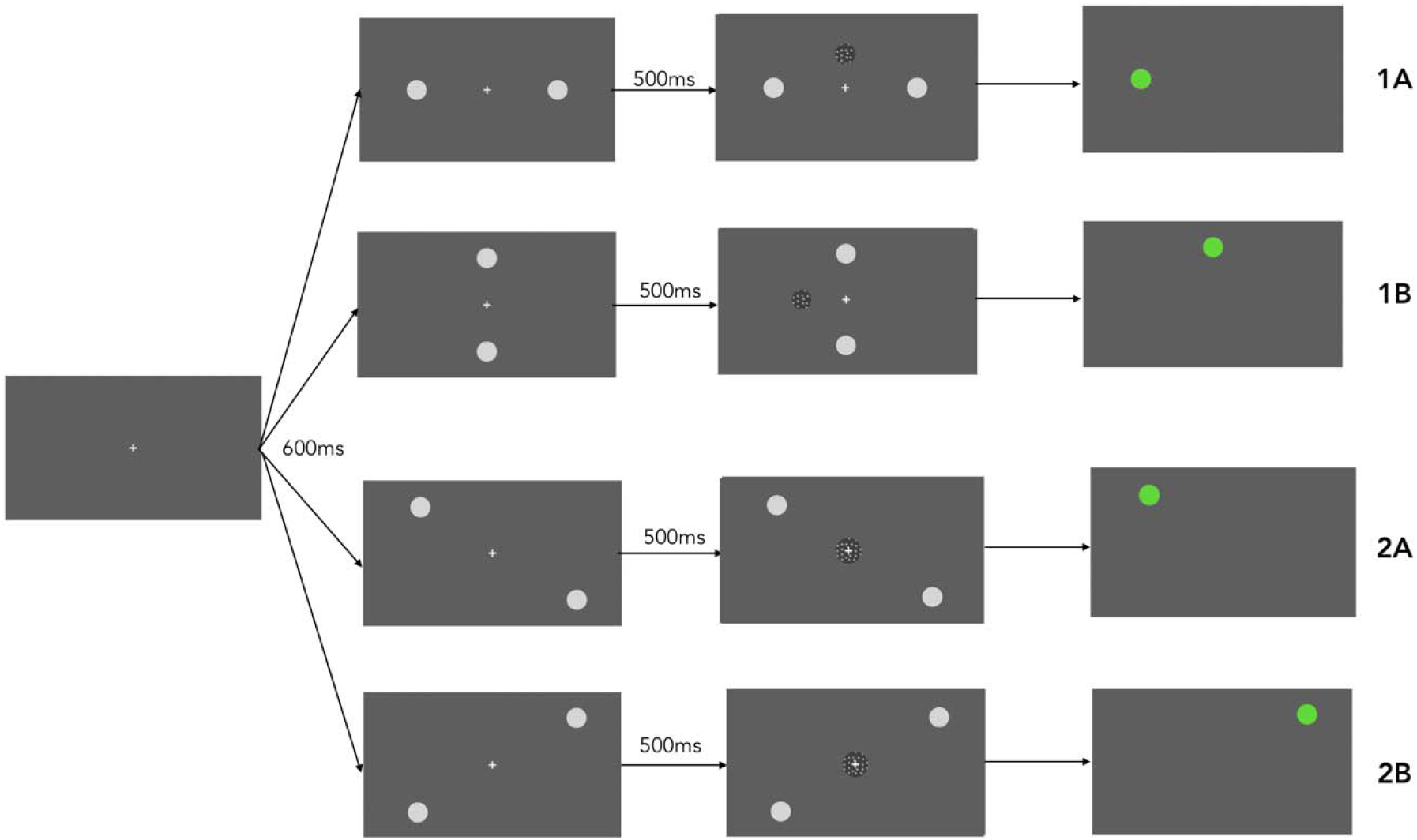
Task Structure and Protocol. A schematic representation of the experimental design. Participants completed four training sessions followed by a single test session. The training sessions were conducted on the same day (Experiment 1 and 2) or across five consecutive days (Experiment 3). Each session included four free-response blocks and one speeded-decision block.

Participants signaled their readiness to begin by pressing the space bar. When participants maintained their gaze on the fixation point (FP, within a 3° by 3° window) for 600ms, two choice targets (RGB = 125, diameter = 1°) would appear. After a delay of 500ms, RDM was shown to the participants. The location of the targets, the location of the RDM, and the direction of the RDM were different between tasks and will be explained fully for each experiment (please see below). RDM was displayed until participants made a saccade (within a 3° by 3° window) to one of the two targets (within 100ms after eyes left the FP). After the saccade, the RDM and the non-chosen target disappeared. The color of the chosen target indicated the outcome of the response—green for correct, red for incorrect.

### Experiment 1A

In this experiment, the targets appeared on the horizontal meridian (7° eccentricity) to the right and left of the FP. The direction of dots of the RDM was along the horizontal axis (i.e., towards right or left). For half of the participants, RDM appeared at the top of the FP (5°) in the training sessions and below the FP (5°) in the test session. For the other half of the participants, the location of the RDM was opposite (below the FP in the training sessions and above the FP in the test session).

### Experiment 1B

The location of the targets was on the vertical meridian (7° eccentricity), above and below the FP. The direction of dots of the RDM was along the vertical axis (i.e., towards up or down). For half of the participants, RDM appeared left of the FP (5°) in the training sessions and right of the FP (5°) in the test session. For the other half of the participants, the location of the RDM was opposite (right of the FP in the training sessions and left of the FP in the test session).

### Experiment 2A

The location of the targets was on the main diagonal (i.e., top-left and bottom right, 7° eccentricity). RDM appeared at the FP location. For half of the participants, the direction of the dots was along the horizontal axis during the training sessions and along the vertical axis on the test session. For the other half of the participants, the direction of the RDM was opposite (vertical axis on training sessions and horizontal axis on the test session).

### Experiment 2B

The location of the targets was on the anti-diagonal (i.e., top-right and bottom left, 7° eccentricity). RDM appeared at the FP location. For half of the participants, the direction of the dots was along the horizontal axis on the training sessions and along the vertical axis on the test session. For the other half of the participants, the direction of the RDM was opposite (vertical axis on training sessions and horizontal axis on the test session).

### Experiment 3

In experiments 1 and 2, the training and test sessions occurred on one day. To investigate the effects of sleep and spacing on decision making, we ran Experiment 3, in which we enrolled participants to perform one session per day (a total of 5 days). The design of Experiment 3 was the same as in Experiment 1B in other regards.

### Data collection

We recruited 34 participants (32 naïve participants and the two authors) with normal or corrected-to-normal vision. Each participant was enrolled in only one experiment: Experiment 1A = 7, Experiment 1B = 7, Experiment 2A = 7, Experiment 2B = 7, Experiment 3 = 6 (including 2 authors). The study was approved by the Ethics Committee at the School of Cognitive Science, IPM, and written informed consent was given by the participants before the start of the study.

### Data analysis

First, we calculated the accuracy (excluding the 0% coherence) and the mean reaction time (for correct response in non-zero coherence, and all trials in 0 coherence trials) of each subject on each session in the Free response (FR) trials.

Then, we used the data from FR trials to fit a drift diffusion model with collapsing bounds and leaky integration using the PyDDM Python package (Shinn et al., 2020) for each session and each participant.

The predicted choice (P) and mean RT of the closed-form classical DDM are described by:

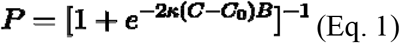

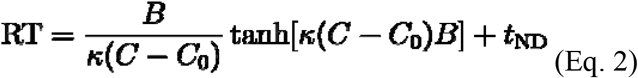

where C is the signed stimulus coherence, C_0_ is a bias term, κ is the drift rate, and B is the boundary. For each session and each subject, the non-decision time (t_ND_) was calculated as the mean of the reaction time in the SD trials. In a separate analysis we dispensed with SD trials and inferred t_ND_ as a fitted variable. The results of these two approaches were qualitatively the same.

We added collapsing bounds and leaky integration to the classical model. These additions mean that the generalized DDM (GDDM) can now only be described in a differential form:

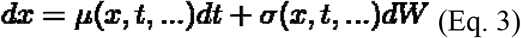

Where μ is the instantaneous drift rate, σ is the instantaneous noise, and W is the Wiener process.

PyDDM numerically solves the GDDM using the Fokker-Planck equation (Shinn et al., 2020). To add drift leak and collapsing bounds, μ and B are described as:

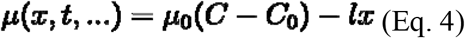

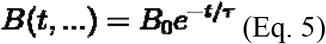

Here ***μ***_**0**_ is the drift rate, ***ι*** is the leak constant, ***B***_**0**_ is the initial boundary height, and τ is the boundary collapse rate.

The model parameters were concurrently fitted by maximizing the log-likelihood of the observed RT on each trial given the model parameters, stimulus strength, and choice. We ended up with five fitted parameters of drift rate **(*μ***_**0**_), drift leak **(*ι*)**, boundary height **(*B***_**0**_), boundary collapse rate **(*τ*)**, and bias (C_0_) for each session and each subject.

To confirm the robustness of our results, we fitted two alternative models to the data as well: (1) a classical DDM (without neither leak nor boundary collapse, i.e., two parameters), (2) two alternative 3-parameter GDDMs (without either leak or boundary collapse). The observed changes in these models and a function of training or transfer were qualitatively similar to the full GDDM.

We calculated a learning index (LI) for the accuracy, RT, ***μ***_**0**_, **ι, *B***_**0**_, and ***τ***, separately. LI was defined:

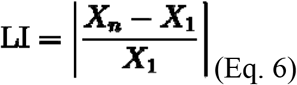

Where X_n_ is the value on the last training session and X_1_ is the value on the first training session (Fine & Jacobs, 2002). The larger the LI, the stronger the learning relative to the first day.

We also calculated a specificity index (SI) as:

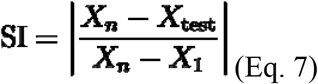

Where X_n_ is the value on the last training session, X_1_ is the value on the first training session, and X_test_ is the value on the test session (Ahissar & Hochstein, 1997). If the learning accrued from X_1_ to X_n_ is not transferred at all, then X_test_ reverts back to X_1_, and the SI will be 1. If the learning is fully transferred from the last training session (X_n_) to the test session (X_test_), SI will be 0.

To check the appropriateness of pooling data across experiments, we fit linear mixed-effects models and tested the statistical significance of experiment type on the regressands (accuracy, RT, and fitted parameter values). The statistical significance of all observed trends (accuracy, RT, µ_0_, □, B_0_, and τ) was assessed using the Page L trend test, a non-parametric test for ordered alternatives in repeated-measures designs (Page, 1963). The test was applied separately to each experiment group, with session number treated as an ordinal predictor. When a significant trend was observed, we conducted post hoc pairwise comparisons between adjacent sessions using pairwise Student’s t-tests with Bonferroni correction. We also tested the statistical significance of between-parameter differences in LIs and SIs using pairwise Student’s t-tests.

## Results

### 1. Accuracy improved and reaction time decreased as a function of training

Across all experiments, participants demonstrated robust improvements in accuracy (**Figure 2A**) and a decrease in reaction time (**Figure 2B**) over the course of the four-session training phase. The fitted linear mixed-effects model showed no effect of experiment type on accuracy (p = 0.41) or RT (p=0.17). Accuracy increased gradually from session 1 (61.3 ± 5.2%), to session 2 (64.4 ± 4.7%), session 3 (68.1 ± 6.2%), and session 4 (74.6± 4.5%, mean ± SEM, across subjects). A Page L trend test showed the significant effects of the training on accuracy (L = 2438, p < 0.001). Post hoc comparisons (paired t-test, Bonferroni correction) revealed no significant difference between sessions 1 and 2 (p = 0.19), session 2 and 3 (p = 0.29), or session 3 and 4 (p = 0.12). We observed a statistically significant improvement in performance from session 1 to 3 (p = 0.032), session 2 to 4 (p = 0.042), and session 1 to 4 (p < 0.001).

**Figure 2:**
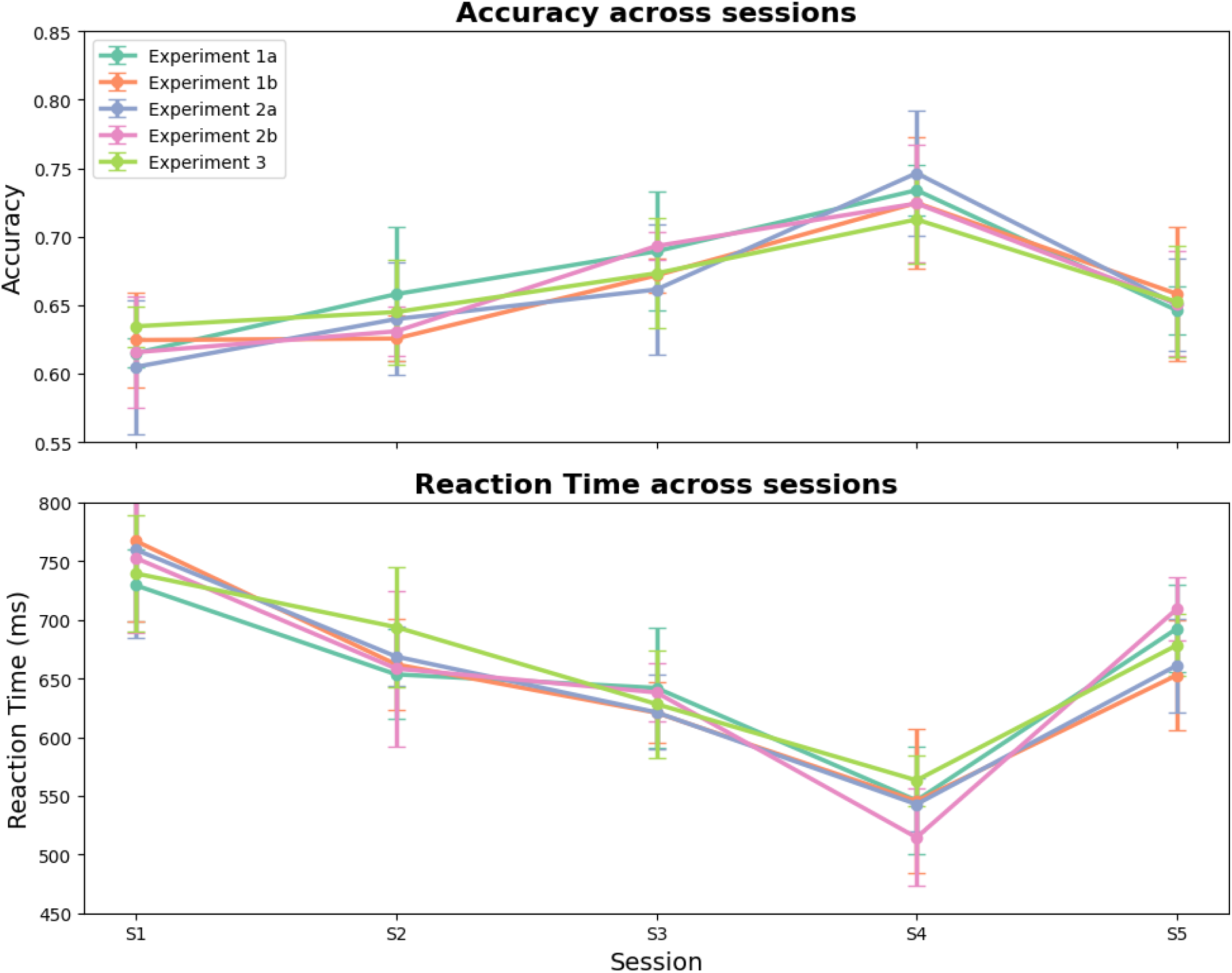
Accuracy and Reaction Time as a Function of Training. **(A)**Accuracy improved steadily across the four training sessions and decreased on the test session. Performance on the test session was significantly lower than on the last training session but remained higher than on the first training session. **(B)**Reaction times (RT) decreased across the training sessions and increased on the test session. RT on the test session was significantly longer than on the last training session, but still shorter than on the first session.

Reaction times (RTs) showed a complementary trend, decreasing across training (session 1 = 748 ± 98 ms, session 2 = 672 ± 57 ms, session 3 = 642 ± 48 ms, session 4 = 549 ± 81 ms; **Figure 2B**). The fitted linear mixed-effects model showed no effect of experiment type on RT (p = 0.17). The reduction in RT was statistically significant (L = 2101, p < 0.001), with no significant change between sessions 1 and 2 (p = 0.18), session 2 and 3 (p = 0.31), or session 3 and 4 (p = 0.07). We observed a significant decrease in RT from session 1 to 3 (p = 0.044), from session 2 to 4 (p = 0.021), and from session 1 to 4 (p < 0.001). This pattern suggests that participants not only became more accurate over training, but also faster in making a decision. In other words, there is no speed-accuracy tradeoff.

On the fifth session (the test session), participants were tested with the RDM at a different location (Experiment 1) or RDM with an orthogonal axis of motion (Experiment 2). Accuracy on the test session (67.7 ± 3.2%) decreased significantly relative to session 4 (74.6 ± 4.5%, paired t-test, t = 5.33, p < 0.001), but it was larger than the performance on session 1 (61.3 ± 5.2%, t = 1.76, p = 0.043). Similarly, RT on the test session (682 ± 45 ms) was significantly longer than RT on session 4 (549 ± 81 ms, t = 4.81, p < 0.001), but it was shorter than RT in session 1 (t = 2.31, p = 0.014). So, learning partially transfers to new locations (experiment 1) or new directions (experiment 2), as performance and RT deteriorate in the test session (relative to session 4), but the values are still better than session 1.

To quantify learning dynamics, we computed a learning index (LI) for each participant, defined as the difference in the values between the last training day and the first day, divided by the value on the first day (Fine & Jacobs, 2002). The mean LI across all subjects was 0.89 ± 0.21 for accuracy and 0.93 ± 0.31 for RTs (mean ± SEM), confirming reliable gains in performance and RT. These training gains establish a baseline for assessing transfer and generalization on the test day. To quantify the extent of transfer and generalization, we computed a specificity index (SI) for both accuracy and RT, as the difference between values on last training day and test day divided by the difference between values on the last training day and the first day (Ahissar & Hochstein, 1997). The mean SI was 0.56 ± 0.04 for accuracy and 0.58 ± 0.05 for RT (mean ± SEM), indicating that roughly half of the training benefit was retained under novel stimulus conditions.

Taken together, these findings support the conclusion that training-induced improvements in perceptual decision making generalized partially, but not fully, to a new stimulus condition. This partial transfer may reflect the coexistence of both stimulus-dependent and stimulus-independent (or decision-related) learning components, a hypothesis we explored further using model-based analysis.

### 2. GDDM Parameters showed unique patterns of change as a function of training

To investigate the cognitive mechanisms underlying behavioral improvements and their generalization, we fitted a generalized drift diffusion model (GDDM) to each participant’s accuracy and RT from the free response trials for each session, allowing us to track changes across training and test sessions. To have a more accurate understanding, we calculated the non-decision time from the speeded decision trials and used that in the GDDM as a fixed parameter. GDDM explained both psychometric and chronometric functions well **(Figure 3)**. The model provided an acceptable fit to the data for each individual subject and session. The goodness-of-fit was consistently high, with all of the coefficient of determination (R^2^) observed across all subjects and sessions were above 0.83. As can be seen in Figure 3, psychometric functions steepened from session 1 to session 4, and flattened on the test session. Chronometric functions shifted downwards vertically from session 1 to 4, and then moved upwards on the test session. We investigated drift rate (***μ***_**0**_), boundary height (***B***_**0**_), drift leak (***l***), and boundary collapse rate (***τ***) as a function of learning and the changed stimulus (Figure 4).

**Figure 3:**
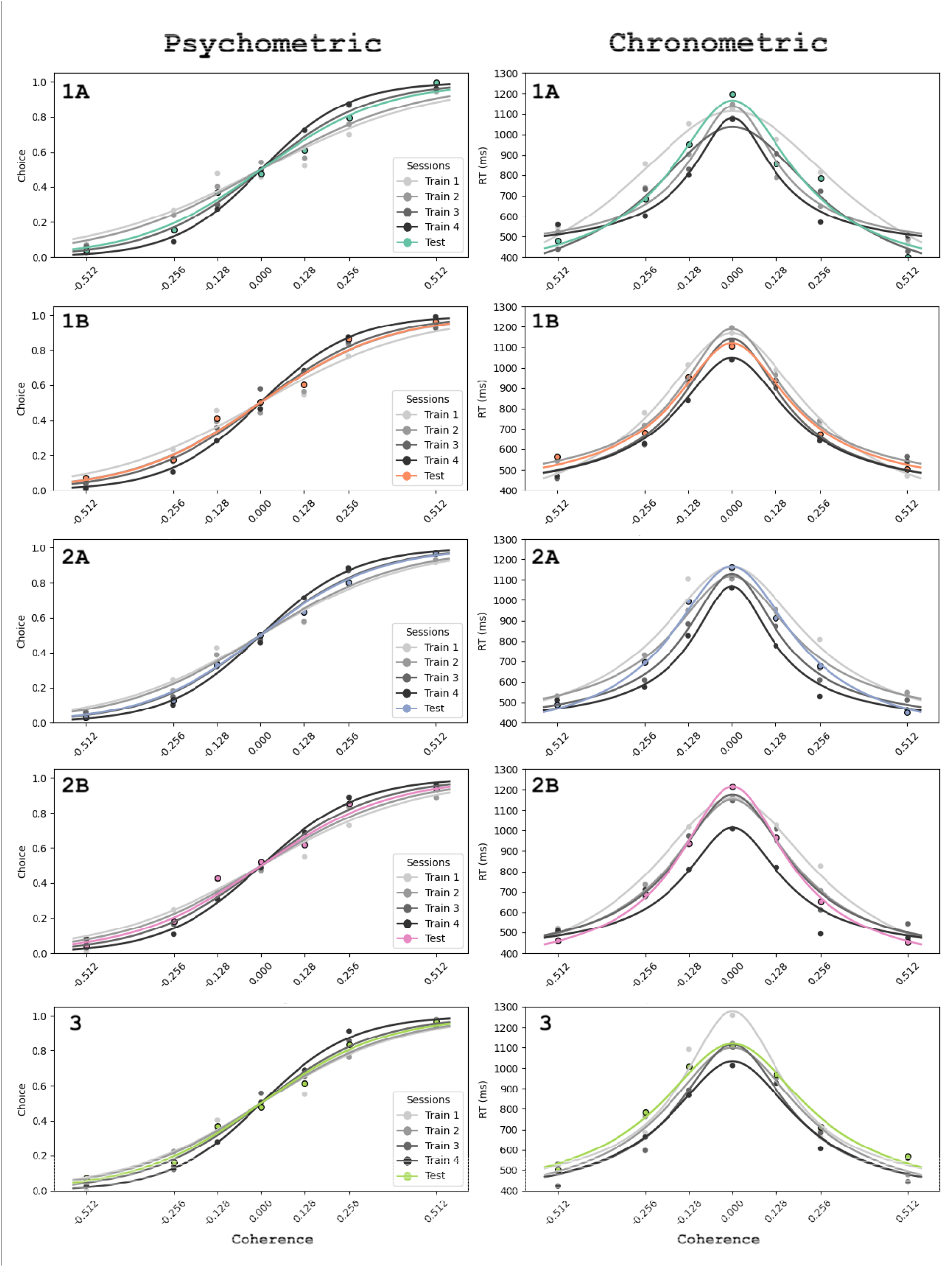
Psychometric and Chronometric Functions. Psychometric (left column) and chronometric (right column) functions fitted (from GDDM, lines) to the averaged data (circle) are shown for each session (for presentation purpose only). Psychometric functions, which plot accuracy against coherence, steepened from session 1 to session 4 and flattened on the test session. Chronometric functions, which plot reaction time against coherence, shifted vertically downwards from session 1 to session 4 and moved upwards on the test session.

**Figure 4:**
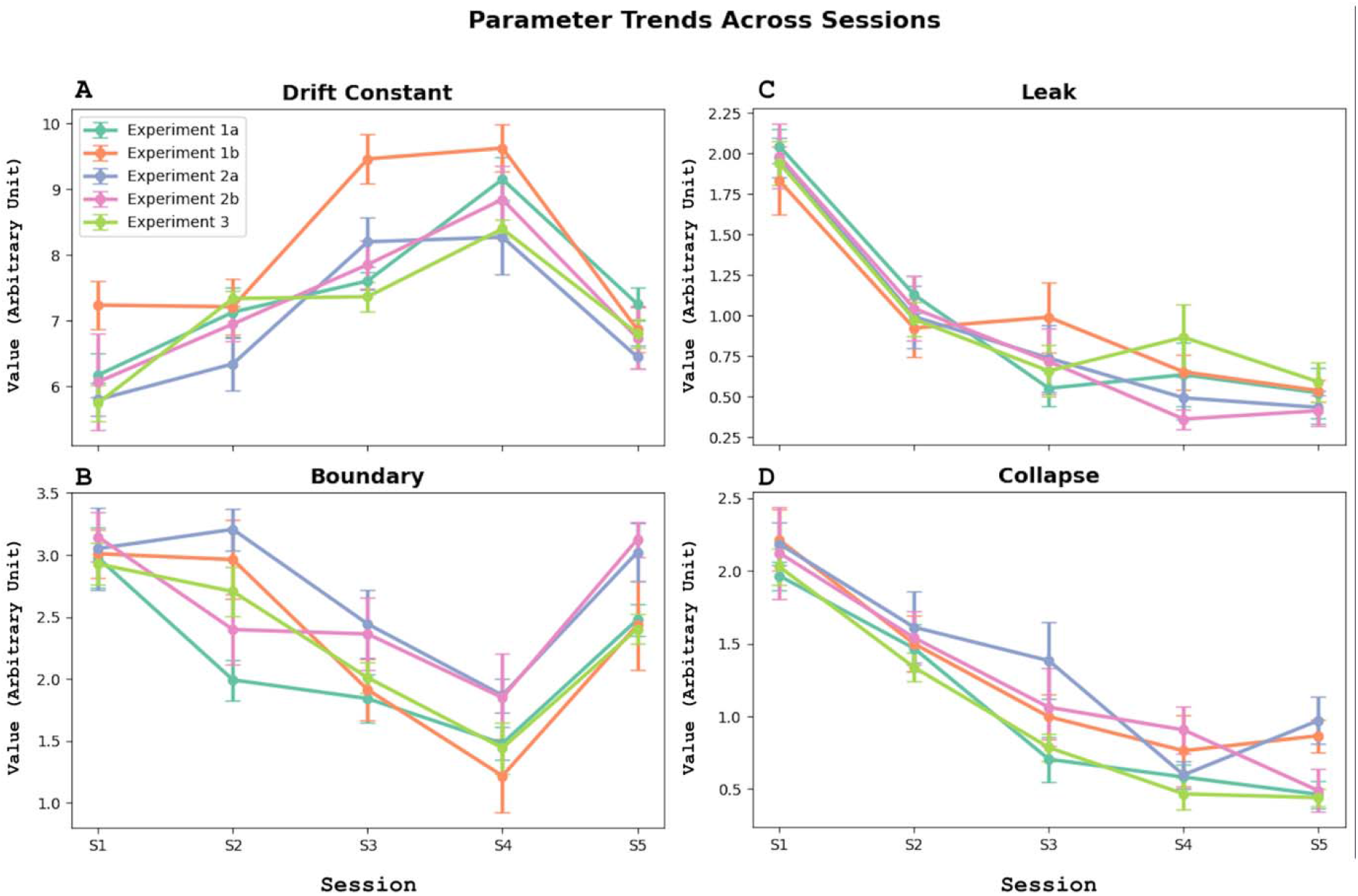
Generalized Drift Diffusion Model (GDDM) Parameter Trends. The trends of the fitted GDDM parameters are shown as a function of training and stimulus change. The drift rate (A) and boundary height (B) increased and decreased, respectively, during training and partially reverted to their pre-training values on the test session. The drift leak (C) and boundary (D) collapse rate significantly decreased over training and showed no significant change on the test session.

We found that **drift rate** (***μ***_**0**_) increased from session 1 (6.49 ± 0.62) to session 2 (6.94 ± 0.49), session 3 (8.21 ± 1.13), and session 4 (8.72 ± 1.22) (Figure 4A). This improvement was statistically significant (Page Test, L = 3122, p = 0.0014). Our fitted linear fixed-effects model showed a significant interaction between experiment type and ***μ***_**0**_ (p = 0.049). Post hoc comparisons revealed no significant difference between sessions 1 and 2 (p =0.41), session 2 and 3 (p = 0.063), or session 3 and 4 (p = 0.28). We observed a significant improvement in drift rate from session 1 to session 3 (p = 0.013), from session 2 to session 4 (p = 0.023), and from session 1 to session 4 (p < 0.001). The drift rate on the test session (7.23 ± 0.42) decreased significantly relative to session 4 (paired t-test, t = 4.18, p < 0.001), but it was larger than the drift rate on session 1 (t = 3.03, p = 0.0027). Learning index (LI, Figure 5A, green column) and specificity index (SI, Figure 5B, green column) for drift rate were 0.49 ± 0.09 and 0.62 ± 0.102, respectively.

**Figure 5:**
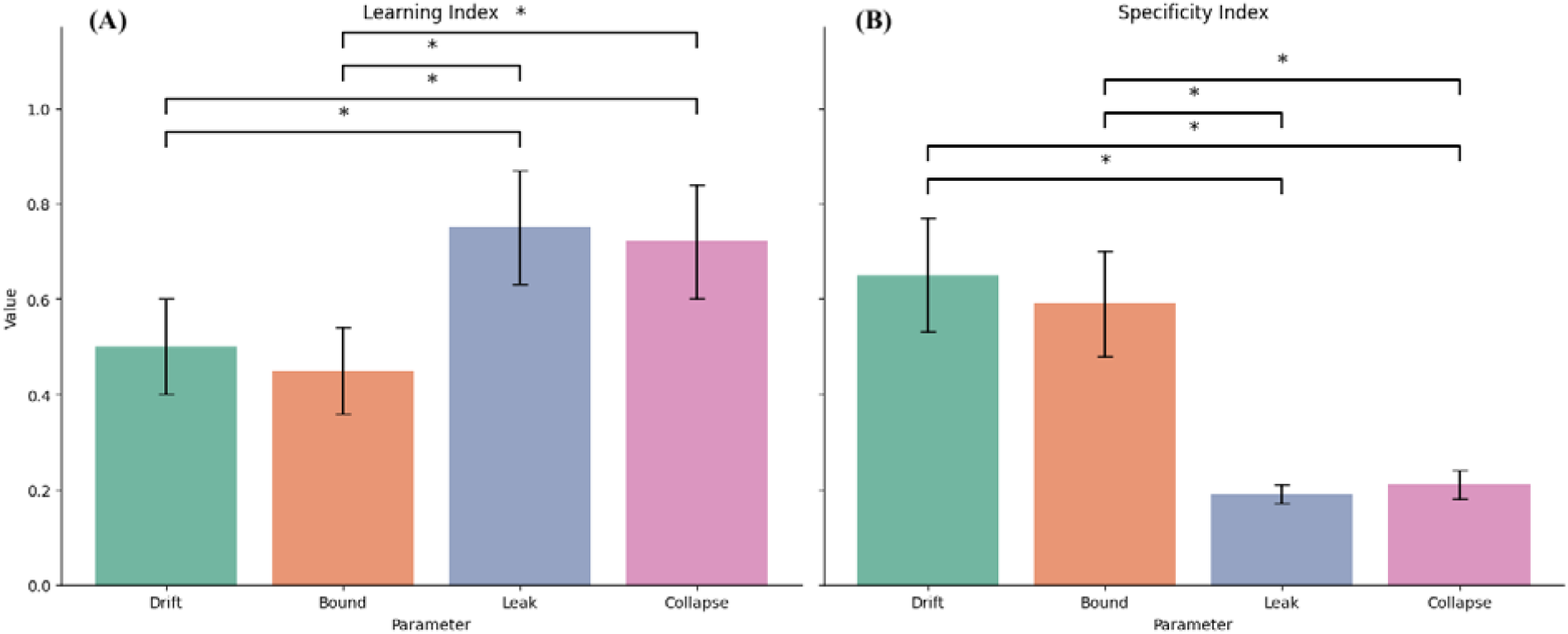
Quantifying Learnability and Specificity of GDDM Parameter Changes. **(A)**Learning Index (LI): leak (blue) and boundary collapse (pink) rate exhibited higher LI values than drift (green) or boundary height (orange), indicating more significant learning-related changes compared to drift rate and boundary height. * p < 0.05. **(B)**Specificity Index (SI): leak (blue) and boundary collapse (pink) rate showed a lower SI, indicating greater generalization to new stimulus features, in contrast to drift rate (green) and boundary height (orange), which had higher SI values and less generalization. * p < 0.05.

For the **boundary height (μ**_**0**_), we observed a significant decrease from session 1 (3.08 ± 0.47) to session 2 (2.69 ± 0.83), session 3 (2.34 ± 0.52) to session 4 (1.53 ± 0.53, L = 2853, p =0.0032) (Figure 4B). The fitted linear fixed-effects model showed no significant interaction between experiment type and boundary height (p = 0.11). Post hoc comparisons revealed no significant difference between sessions 1 and 2 (p = 0.53), session 2 and 3 (p = 0.32), or session 3 and 4 (p = 0.17). We observed a significant decrease in boundary height from session 1 to session 3 (p = 0.031), from session 2 to session 4 (p = 0.026), and from session 1 to session 4 (p < 0.001). The boundary height on the test session (2.58 ± 0.56) increased significantly relative to session 4 (paired t-test, t = 4.18, p < 0.001), but it was larger than the boundary height on session 1 (t = 2.36, p = 0.0129). LI (Figure 5A, orange column), and SI Figure 5B, orange column) for boundary height were 0.41 ± 0.07 and 0.59 ± 0.098, respectively.

The **drift leak (*l*)** showed a significantly decreasing trend from session 1 (1.81 ± 0.18) to session 2 (1.08 ± 0.13), session 3 (0.89 ± 0.23), and session 4 (0.66 ± 0.10, L = 4021, p < 0.001) (Figure 4C). The fitted linear fixed-effects model showed no significant interaction between experiment type and drift leak (p = 0.23). Post hoc comparisons revealed a significant difference between sessions 1 and 2 (p < 0.001), sessions 1 and 3 (p < 0.001), and sessions 2 and 4 (p = 0.032), but no differences were observed between sessions 2 and 3 (p = 0.44) or sessions 3 and 4 (p = 0.39). Importantly, we did not see a significant difference in the drift leak between session 4 and the test session (0.64 ± 0.08, t = 0.73, p = 0.24). The drift leak on the test session was significantly lower than on session 1 (t = 4.26, p < 0.001). The LI (Figure 5A, blue column) for the drift leak was 0.73 ± 0.13, and the SI (Figure 5B, blue column) was 0.19 ± 0.03.

The **boundary collapse rate (τ)** showed a similar trend to the drift leak. The linear fixed-effects model showed no significant interaction between experiment type and boundary collapse rate (p = 0.17). There was a statistically significant decrease from session 1 (2.21 ± 0.31) to session 2 (1.71 ± 0.28), session 3 (1.38 ± 0.64), and session 4 (0.83 ± 0.24, L = 3119, p < 0.001). Post hoc comparison showed no statistically significant differences between session 1 and 2 (p = 0.072), session 2 and 3 (p = 0.121), or session 3 and 4 (p = 0.183). There was a significant decrease from session 1 to 3 (p = 0.024), 2 to 4 (p = 0.014), and 1 to 4 (p < 0.001). Similar to the situation we observed with the drift leak, and in contrast to drift rate and boundary height, there was no statistically significant change in the boundary collapse rate on the test session (0.78 ± 0.31) and session 4 (t = 0.42, p = 0.34). This value was significantly lower than the one observed on session 1 (t = 5.25, p < 0.001). The LI (Figure 5A, pink column) for the boundary collapse rate was 0.71 ± 0.13, and the SI (Figure 5B, pink column) was 0.21 ± 0.03.

### 3. Quantifying and comparing the learnability and specificity of GDDM parameters

GDDM parameters showed distinct patterns as a function of learning and transfer; drift rate and boundary height partially reverted to pre-training values on the test day, while leak and collapse parameters remained stable. To quantify the differences in learnability and transferability (stimulus specificity) between these parameters, we applied pairwise Student’s t-tests comparing the values of the learning index (LI) and specificity index (SI).

For LI, Drift leak (***l***) and boundary collapse rate (***τ***) both exhibited more training-related changes (as shown by higher LIs (0.73□± □0.13 and 0.71□± □0.13, respectively), compared to the LI for drift rate (µ, 0.49□± □0.09) and the boundary height (B_0_, 0.41□± □0.07), that showed more modest learning-related changes **(Figure 5A)**. While there was no difference between drift leak and boundary collapse rate (t = 0.18, p = 0.42), or between drift rate and boundary height (t = 0.46, p = 0.32), there was a significant difference between drift leak and drift rate (t = 2.31, p < 0.02), between drift leak and boundary height (t = 2.42, p < 0.02), between boundary collapse rate and drift rate (t = 2.18, p < 0.02), and between boundary collapse rate and boundary height (t = 2.37, p < 0.02).

For SI, Drift leak (***l***) and boundary collapse rate (***τ***) both exhibited more generalization to new stimulus features, as shown by SI closer to 0 (0.19□± □0.03 and 0.21□± □0.03, respectively), compared to the SI for drift rate (0.62□± □0.10) and the boundary height (0.59□± □0.09), that showed less generalization **(Figure 5B)**. While there was no difference between drift leak and boundary collapse rate (t = 0.31, p = 0.38), or between drift rate and boundary height (t = 0.98, p = 0.17), there was a significant difference between drift leak and drift rate (t = 3.02, p < 0.003), between drift leak and boundary height (t = 2.80, p < 0.006), between boundary collapse rate and drift rate (t = 2.93, p < 0.004), and between boundary collapse rate and boundary height (t = 2.77, p < 0.006).

## Discussion

We investigated how perceptual learning alters decision dynamics in a direction-discrimination task, and how such improvements generalize to untrained stimulus parameters. We found that participants showed improvement in both accuracy and reaction time across four training sessions. On a fifth test session, when the stimulus location or direction of motion (in separate participants) was changed, performance declined relative to the last day of training, but still better than the first day, suggesting partial generalization. Model-based analysis using a generalized drift diffusion model (GDDM) revealed that learning was associated with an increase in drift rate, and a decrease in drift leak, boundary height, and boundary collapse rate. Comparing these values on the test session with the last training session showed that while drift rate and boundary height were partially generalized to new stimulus features, drift leak and boundary collapse rate were fully generalized.

The observed difference in generalization between different parameters supports the hypothesis that perceptual learning comprises multiple, separable components, each governed by different mechanisms (Ahissar & Hochstein, 2004). Classic perceptual learning studies emphasized the specificity of perceptual learning to low-level stimulus features such as orientation (Fahle & Edelman, 1993), location (Fahle & Morgan, 1996), direction of motion (Ball & Sekuler, 1982), or spatial frequency (Fiorentini & Berardi, 1980), leading to the view that changes occur primarily in early visual areas (Carmel & Carrasco, 2008; Schoups et al., 2001). However, subsequent studies have shown that perceptual improvements can generalize across space, modality, and task structure under certain conditions (Green et al., 2010; Xiao et al., 2008), implying a role for higher-order cognitive processes.

Electrophysiological work in non-human primates adds nuance to this debate. For example, Law and Gold reported that in motion discrimination tasks, learning was associated with changes in decision-related areas in the parietal cortex, but not in the middle temporal (MT), despite MT being the canonical motion-processing area (Law & Gold, 2008). Similarly, Yang & Maunsell found only weak or feedback-driven effects in V1, contrasting with expectations of strong plasticity in early visual cortex (Yang & Maunsell, 2004, Sanayei et al., 2018). These findings suggest that stimulus-evoked changes may be too subtle or distributed for standard single-unit recordings to detect. Human fMRI studies, by contrast, have reported both decision-level changes in frontoparietal circuits and subtle tuning changes in motion-sensitive extrastriate areas (e.g., hMT+/V5) (Chen et al., 2016; Shibata et al., 2016). Differences in training paradigms may partly explain the discrepancy: monkey studies often varied stimulus location and direction daily due to receptive field constraints, whereas human psychophysics and fMRI studies typically used fixed stimulus configurations. Psychophysical paradigms such as ‘roving’ or ‘double training’ further show that transfer can be promoted or abolished depending on feature variability during practice (Dosher et al., 2020; Xiao et al., 2008; Chen et al., 2013). Together, these cross-species and methodological differences highlight that the locus of plasticity in perceptual learning is sensitive to both neural recording scale and training regimen.

Our results provide new evidence in support of the view that perceptual learning comprises multiple, separable components. Improvements in drift leak and boundary collapse time constants during training, parameters associated with how evidence is accumulated and decision criteria are stabilized, suggest that participants adopted more efficient decision policies over training. These parameters remained stable even when the stimulus location or direction of motion was changed, indicating that decision-level optimizations are not tied to specific sensory features. This aligns with findings from neurophysiological studies in monkeys, which have shown that training-related changes in perceptual decision-making often emerge in the parietal rather than early visual cortex (Law & Gold, 2008), and with human neuroimaging data implicating changes in frontoparietal circuits as a function of training (Jia et al., 2018; Shibata et al., 2016).

In contrast, drift rate and boundary height, parameters more directly associated with the quality and magnitude of extracted sensory evidence, deteriorated when stimulus location or axis of motion changed. This suggests that these components of the learned behavior were specific to the trained stimulus configuration. Our findings thus extend earlier work using standard DDMs (Petrov et al., 2011) by demonstrating that more complex, time-resolved models like the GDDM can parse learning into fully-generalizable and partially-generalizable components. The dissociation supports hybrid models of perceptual learning, where early-stage sensory tuning is accompanied by higher-level reweighting or reinterpretation of sensory signals (Ahissar & Hochstein, 2004; Dosher & Lu, 1998).

The use of GDDMs in this context is novel and justified. Most previous work has focused on drift rate, boundary height, and non-decision time (Gold et al., 2010), but the current model’s inclusion of leaky integration and collapsing bounds allowed us to capture changes in decision strategies (Stine et al., 2020). These components are particularly relevant in free-response tasks, where subjects must manage speed–accuracy tradeoffs. Our data suggest that improvements in leak and boundary stability are reliable indicators of strategy-level learning and should be more widely measured in future work.

The absence of full generalization also constrains models of perceptual plasticity. If decision-level parameters alone were sufficient for transfer, we would expect little to no performance loss on the test day. Instead, the decline in accuracy (and accompanying decrease in drift rate and the increase in bound height) suggests that stimulus-specific representations, perhaps in early visual areas, continue to limit transfer. This underscores the layered architecture of perceptual learning, in which strategy changes must be supported by stable sensory input to drive robust generalization.

A further question concerns the role of sleep-dependent consolidation in perceptual learning. Previous studies using texture or orientation discrimination tasks have reported that performance improvements often emerge or stabilize following sleep, suggesting that offline consolidation processes contribute critically to visual plasticity (Gais et al., 2000; Karni et al., 1994; Stickgold et al., 2000). In our experiment, a separate group of participants completed the training and test sessions on consecutive days, allowing us to probe whether sleep facilitated either learning or transfer. However, we found no significant differences between the sleep and same-day training groups in terms of improvements in accuracy, response speed, or any of the GDDM parameters. This null result may indicate that consolidation is not always critical for at least some forms of perceptual learning, specifically motion discrimination learning. However, this inference is limited by the small number of participants in our sleep cohort.

Several future directions follow from this work. First, given that decision-level parameters like drift leak generalized even when stimulus-level performance declined, it would be valuable to explore how explicit training on decision strategies, e.g., via feedback manipulation or metacognitive cues, can accelerate learning or enhance transfer. Second, this modeling framework can be extended to other modalities and clinical populations, such as aging individuals or patients with perceptual deficits, where isolating decision dynamics could inform cognitive remediation strategies. Finally, future work combining this modeling approach with neurophysiological or neuroimaging data will be necessary to validate the underlying mechanisms.

More broadly, our results suggest that perceptual learning is not simply a matter of tuning early sensory filters, but also involves optimizing how evidence is accumulated and acted upon over time. By applying principled cognitive modeling, we can begin to deconstruct the layered architecture of experience-dependent learning and ultimately, use this knowledge to guide adaptive learning systems both in natural and artificial intelligence (Cheng et al., 2025).

## References

Ahissar, M., & Hochstein, S. (1997). Task difficulty and the specificity of perceptual learning. Nature, 387(6631), 401–406. 10.1038/387401a0

Ahissar, M., & Hochstein, S. (2004). The reverse hierarchy theory of visual perceptual learning. Trends in Cognitive Sciences, 8(10), 457–464. 10.1016/j.tics.2004.08.011

Ball, K., & Sekuler, R. (1982). A Specific and Enduring Improvement in Visual Motion Discrimination. Science, 218(4573), 697–698. 10.1126/science.7134968

Brainard, D. H. (1997). The Psychophysics Toolbox. Spatial Vision, 10(4), 433–436.

Carmel, D., & Carrasco, M. (2008). Perceptual Learning and Dynamic Changes in Primary Visual Cortex. Neuron, 57(6), 799–801. 10.1016/j.neuron.2008.03.009

Chen, N., Bi, T., Zhou, T., Li, S., Liu, Z., & Fang, F. (2015). Sharpened cortical tuning and enhanced cortico-cortical communication contribute to the long-term neural mechanisms of visual motion perceptual learning. NeuroImage, 115, 17–29. 10.1016/j.neuroimage.2015.04.041

Chen, N., Cai, P., Zhou, T., Thompson, B., & Fang, F. (2016). Perceptual learning modifies the functional specializations of visual cortical areas. Proceedings of the National Academy of Sciences, 113(20), 5724–5729. 10.1073/pnas.1524160113

Chen, X., Sanayei, M., & Thiele, A. (2013). Perceptual learning of contrast discrimination in macaca mulatta. Journal of Vision, 13(13), 22–22. 10.1167/13.13.22

Cheng, Y.-A., Sanayei, M., Chen, X., Jia, K., Li, S., Fang, F., Watanabe, T., Thiele, A., & Zhang, R.-Y. (2025). A neural geometry approach comprehensively explains apparently conflicting models of visual perceptual learning. Nature Human Behaviour, 9(5), 1023–1040. 10.1038/s41562-025-02149-x

Dosher, B. A., Liu, J., Chu, W., & Lu, Z.-L. (2020). Roving: The causes of interference and re-enabled learning in multi-task visual training. Journal of Vision, 20(6), 9. 10.1167/jov.20.6.9

Dosher, B. A., & Lu, Z.-L. (1998). Perceptual learning reflects external noise filtering and internal noise reduction through channel reweighting. Proceedings of the National Academy of Sciences, 95(23), 13988–13993. 10.1073/pnas.95.23.13988

Doyon, J., Korman, M., Morin, A., Dostie, V., Tahar, A. H., Benali, H., Karni, A., Ungerleider, L. G., & Carrier, J. (2009). Contribution of night and day sleep vs. Simple passage of time to the consolidation of motor sequence and visuomotor adaptation learning. Experimental Brain Research, 195(1), 15–26. 10.1007/s00221-009-1748-y

Fahle, M., & Edelman, S. (1993). Long-term learning in vernier acuity: Effects of stimulus orientation, range and of feedback. Vision Research, 33(3), 397–412. 10.1016/0042-6989(93)90094-D

Fahle, M., & Morgan, M. (1996). No transfer of perceptual learning between similar stimuli in the same retinal position. Current Biology, 6(3), 292–297. 10.1016/S0960-9822(02)00479-7

Fine, I., & Jacobs, R. A. (2002). Comparing perceptual learning across tasks: A review. Journal of Vision, 2(2), 5–5. 10.1167/2.2.5

Fiorentini, A., & Berardi, N. (1980). Perceptual learning specific for orientation and spatial frequency. Nature, 287(5777), 43–44. 10.1038/287043a0

Gais, S., Plihal, W., Wagner, U., & Born, J. (2000). Early sleep triggers memory for early visual discrimination skills. Nature Neuroscience, 3(12), 1335–1339. 10.1038/81881

Gold, J. I., Law, C.-T., Connolly, P., & Bennur, S. (2010). Relationships between the threshold and slope of psychometric and neurometric functions during perceptual learning: Implications for neuronal pooling. Journal of Neurophysiology, 103(1), 140–154. 10.1152/jn.00744.2009

Gold, J. I., & Shadlen, M. N. (2007). The Neural Basis of Decision Making. Annual Review of Neuroscience, 30(1), 535–574. 10.1146/annurev.neuro.29.051605.113038

Green, C. S., Pouget, A., & Bavelier, D. (2010). Improved Probabilistic Inference as a General Learning Mechanism with Action Video Games. Current Biology, 20(17), 1573–1579. 10.1016/j.cub.2010.07.040

Hubel, D. H., & Wiesel, T. N. (1962). Receptive fields, binocular interaction and functional architecture in the cat’s visual cortex. The Journal of Physiology, 160(1), 106–154. 10.1113/jphysiol.1962.sp006837

Jia, K., Xue, X., Lee, J.-H., Fang, F., Zhang, J., & Li, S. (2018). Visual perceptual learning modulates decision network in the human brain: The evidence from psychophysics, modeling, and functional magnetic resonance imaging. Journal of Vision, 18(12), 9. 10.1167/18.12.9

Karni, A., Tanne, D., Rubenstein, B. S., Askenasy, J. J. M., & Sagi, D. (1994). Dependence on REM Sleep of Overnight Improvement of a Perceptual Skill. Science, 265(5172), 679–682. 10.1126/science.8036518

Kleiner, M., Brainard, D., Pelli, D., Ingling, A., Murray, R., & Broussard, C. (2007). What’s new in psychtoolbox-3. Perception, 36(14), 1–16.

Law, C.-T., & Gold, J. I. (2008). Neural correlates of perceptual learning in a sensory-motor, but not a sensory, cortical area. Nature Neuroscience, 11(4), 505–513. 10.1038/nn2070

Maertens, M., & Pollmann, S. (2005). fMRI Reveals a Common Neural Substrate of Illusory and Real Contours in V1 after Perceptual Learning. Journal of Cognitive Neuroscience, 17(10), 1553–1564. 10.1162/089892905774597209

Mckee, S. P., & Westhe, G. (1978). Improvement in vernier acuity with practice. Perception & Psychophysics, 24(3), 258–262. 10.3758/BF03206097

Mollon, J. D., & Danilova, M. V. (1996). Three remarks on perceptual learning. Spatial Vision, 10(1), 51–58. 10.1163/156856896X00051

Mukai, I., Kim, D., Fukunaga, M., Japee, S., Marrett, S., & Ungerleider, L. G. (2007). Activations in Visual and Attention-Related Areas Predict and Correlate with the Degree of Perceptual Learning. The Journal of Neuroscience, 27(42), 11401–11411. 10.1523/JNEUROSCI.3002-07.2007

Page, E. B. (1963). Ordered Hypotheses for Multiple Treatments: A Significance Test for Linear Ranks. Journal of the American Statistical Association, 58(301), 216–230. 10.1080/01621459.1963.10500843

Palmer, J., Huk, A. C., & Shadlen, M. N. (2005). The effect of stimulus strength on the speed and accuracy of a perceptual decision. Journal of Vision, 5(5), 1. 10.1167/5.5.1

Pelli, D. G. (1997). The VideoToolbox software for visual psychophysics: Transforming numbers into movies. Spatial Vision, 10(4), 437–442.

Petrov, A. A., Van Horn, N. M., & Ratcliff, R. (2011). Dissociable perceptual-learning mechanisms revealed by diffusion-model analysis. Psychonomic Bulletin & Review, 18(3), 490–497. 10.3758/s13423-011-0079-8

Saarinen, J., & Levi, D. M. (1995). Perceptual learning in vernier acuity: What is learned? Vision Research, 35(4), 519–527. 10.1016/0042-6989(94)00141-8

Sagi, D., & Tanne, D. (1994). Perceptual learning: Learning to see. Current Opinion in Neurobiology, 4(2), 195–199. 10.1016/0959-4388(94)90072-8

Sanayei, M., Chen, X., Chicharro, D., Distler, C., Panzeri, S., & Thiele, A. (2018). Perceptual learning of fine contrast discrimination changes neuronal tuning and population coding in macaque V4. Nature Communications, 9(1). 10.1038/s41467-018-06698-w

Schoups, A., Vogels, R., Qian, N., & Orban, G. (2001). Practising orientation identification improves orientation coding in V1 neurons. Nature, 412(6846), 549–553. 10.1038/35087601

Shibata, K., Sasaki, Y., Kawato, M., & Watanabe, T. (2016). Neuroimaging Evidence for 2 Types of Plasticity in Association with Visual Perceptual Learning. Cerebral Cortex, 26(9), 3681–3689. 10.1093/cercor/bhw176

Shinn, M., Lam, N. H., & Murray, J. D. (2020). A flexible framework for simulating and fitting generalized drift-diffusion models. eLife, 9, e56938. 10.7554/eLife.56938

Stickgold, R., Whidbee, D., Schirmer, B., Patel, V., & Hobson, J. A. (2000). Visual Discrimination Task Improvement: A Multi-Step Process Occurring During Sleep. Journal of Cognitive Neuroscience, 12(2), 246–254. 10.1162/089892900562075

Stine, G. M., Zylberberg, A., Ditterich, J., & Shadlen, M. N. (2020). Differentiating between integration and non-integration strategies in perceptual decision making. eLife, 9, e55365. 10.7554/eLife.55365

Tootell, R., Silverman, M., Hamilton, S., Switkes, E., & De Valois, R. (1988). Functional anatomy of macaque striate cortex. V. Spatial frequency. The Journal of Neuroscience, 8(5), 1610–1624. 10.1523/JNEUROSCI.08-05-01610.1988

Xiao, L.-Q., Zhang, J.-Y., Wang, R., Klein, S. A., Levi, D. M., & Yu, C. (2008). Complete Transfer of Perceptual Learning across Retinal Locations Enabled by Double Training. Current Biology, 18(24), 1922–1926. 10.1016/j.cub.2008.10.030

Yang, T., & Maunsell, J. H. R. (2004). The Effect of Perceptual Learning on Neuronal Responses in Monkey Visual Area V4. The Journal of Neuroscience, 24(7), 1617–1626. 10.1523/JNEUROSCI.4442-03.2004

